# Relationship between footshock intensity, post-training corticosterone release and contextual fear memory specificity over time

**DOI:** 10.1101/643098

**Authors:** Moisés dos Santos Corrêa, Barbara dos Santos Vaz, Gabriel David Vieira Grisanti, Joselisa Péres Queiroz de Paiva, Paula Ayako Tiba, Raquel Vecchio Fornari

**Affiliations:** Centro de Matemática, Computação e Cognição (CMCC), Universidade Federal do ABC (UFABC), São Bernardo do Campo, SP, Brazil; Brain Institute, Hospital Israelita Albert Einstein (HIAE), São Paulo, SP – Brazil

**Keywords:** Corticosterone, fear conditioning, fear memory, memory generalization, memory specificity

## Abstract

Overgeneralized fear has long been implicated in generalized anxiety and post-traumatic stress disorder, however, time-dependent mechanisms underlying memory retrieval are still not completely understood. Previous studies have revealed that stronger fear conditioning training protocols are associated with both increased post-training corticosterone (CORT) levels and fear responses at later retrieval tests. Here we used discriminative contextual fear conditioning (CFC) to investigate the relationship between post-training CORT levels and memory specificity in different retrieval timepoints. Wistar rats were exposed to CFC training with increasing footshock intensities (0.3, 0.6 or 1.0mA) and had their blood collected 30 min afterwards to measure post-training plasma CORT. After 2, 14 or 28 days, rats were tested for memory specificity either in the training or in the novel context. Regression analysis was used to verify linear and non-linear interactions between CORT levels and freezing. Higher footshock intensities increased post-training CORT levels and freezing times during tests in all timepoints. Moreover, stronger trainings elicited faster memory generalization, which was associated with higher CORT levels during memory consolidation. The 0.3mA training maintained memory specificity up to 28 days. Additionally, linear regressions suggest that the shift from specific to generalized memories is underway at 14 days after training. These results are consistent with the hypotheses that stronger training protocols elicit a faster generalization rate, and that this process is associated with increased post-training CORT release.

**HIGHLIGHTS:** Stronger contextual fear conditioning (CFC) elicits higher plasma corticosterone (CORT).

Strong CFC and high CORT levels increase the rate of memory generalization.

Weak CFC and low CORT levels retain memory specificity up to 28 days.

Post-training plasma CORT is linearly associated with remote generalized fear.

## INTRODUCTION

The overgeneralization of fear is associated with generalized anxiety and post-traumatic stress disorder (Dymond et al., 2015; Lissek et al., 2014, 2011) and has recently become a target of intense investigation (Asok et al., 2019; Dymond et al., 2015; Laufer et al., 2016). Contextual fear conditioning (CFC) is a useful and widely recognized task to investigate contextual memory acquisition, consolidation and retrieval. The precision of the memory retrieval (i.e. memory specificity) changes over time and can be evaluated in proper designed CFC protocols, where trained animals are reexposed to either the training context or to a novel context that is slightly different from the original one (Bueno et al., 2017; Fanselow, 1980; Huckleberry et al., 2016; Pedraza et al., 2016; Poulos et al., 2016; Wiltgen et al., 2010; Wiltgen and Silva, 2007; Winocur et al., 2010; Xu and Südhof, 2013). Failure to discriminate between both contexts, as demonstrated by conditioned fear responses, provides a proxy of memory generalization (Hunsaker and Kesner, 2013, 2008; Kesner and Hunsaker, 2010; Rudy and O’Reilly, 1999). At short post-training intervals (1-7 days), fear memories are usually specific to the training context but later become generalized across different contexts (Biedenkapp and Rudy, 2007; de Oliveira Alvares et al., 2012; Haubrich et al., 2016; Wiltgen and Silva, 2007; Winocur et al., 2007). It is suggested that memory becomes more dependent of neocortical regions as time goes by, losing its dependence from the hippocampus. This reallocation is associated with a transformation in the strength and quality of memory, thus, context-dependent memories may transform to semantic-like memories, becoming less detailed and vivid (i.e., generalized) (Asok et al., 2019; Frankland and Bontempi, 2005).

The consolidation of fear memories is modulated by glucocorticoids (GCs) – cortisol in humans and corticosterone (CORT) in rats (De Kloet et al., 1999; de Quervain et al., 2009; Finsterwald and Alberini, 2014; McGaugh and Roozendaal, 2002). Increasing the intensity of the CFC training protocol (i.e. current or number of footshocks) has been shown to elicit higher levels of post-training plasma CORT, which is associated with increased time spent on freezing in retrieval tests (Cordero et al., 1998; Marchand et al., 2007), suggesting that CORT enhances the consolidation of fear memory (Atsak et al., 2012; Cordero and Sandi, 1998; Reul and Kloet, 1985; Roozendaal et al., 2003). It is suggested that CORT acts on both mineralocorticoid receptors (MR) and glucocorticoid receptors (GR) in the hippocampus, amygdala and prefrontal cortex to enhance fear memory (Atsak et al., 2012; Cordero and Sandi, 1998; Reul and Kloet, 1985; Roozendaal et al., 2003). However, the role played by the MR and GR in the process of memory generalization remains unclear.

To date, few studies have explored the relationship between stress hormones and the specificity of contextual fear memories (Atucha et al., 2017; Bueno et al., 2017; Pedraza et al., 2016). Post-training inhibition of plasma CORT levels with metyrapone has been shown to prolong memory specificity (Pedraza et al., 2016). Nevertheless, we have previously reported that the post-training administration of CORT did not affect memory specificity of a single-trial CFC task, in both recent and remote memory tests (Bueno et al., 2017). Hence, the underlying mechanism by which CORT modulates fear memory specificity remains unclear, and we speculate that this process may be affected by the strength of the training protocol and later post-training release of CORT. Here, we investigated how increasing the intensity of footshocks affected post-training endogenous CORT and memory specificity at different retention intervals.

## METHODS

### Subjects

Three-month old male Wistar rats, obtained from *Instituto Nacional de Farmacologia* (INFAR-UNIFESP (total n = 266; weighing between 275-385g at time of training), were kept in controlled conditions of temperature (23 ± 2°C) and light/darkness cycle of 12:12 hours (light phase starting at 7am). Rats were housed in individual cages (20 cm x 16 cm x 18 cm) and provided with food and water *ad libitum*. The rats were adapted to the vivarium for at least one week before the beginning of the experiments. Training and testing were performed during the light phase of the cycle (between 10:00am-03:00pm), at the rat’s nadir of the diurnal rhythm for CORT. All procedures were conducted according to the guidelines and standards of CONCEA - *Conselho Nacional de Controle de Experimentação Animal* (Brazilian Council of Animal Experimentation) and were previously approved by the Ethics Committee on Animal Use - UFABC (CEUA - protocol numbers 5676291015 and 7479070916). Each experiment was conducted with different groups of animals.

### Apparatus

Behavioral experiments were conducted in two identical automated fear conditioning chambers (Med-Associates, Inc., St. Albans, VT), connected to a computer interface for video recording, analysis and measurement of the rat’s freezing behavior in real time. The conditioning box (32 cm wide, 25 cm high and 25 cm deep, VFC-008) was surrounded by a sound attenuating chamber (63.5 cm × 35.5 cm × 76 cm, NIR-022SD) and illuminated by a LED light source (Med Associates NIR-100), which provided visible white light (450-650nm) and invisible near-infrared light (NIR, 940nm) spectra. A NIR video camera (VID-CAM-MONO-4 Fire Wire Video Camera) was attached to the front of the sound attenuating chamber, facing the transparent front wall of the conditioning chamber. The scrambled footshocks were administered through the grid floor of the cage (AC constant current), controlled by an Aversive Stimulator (ENV-414S). A general activity index was derived in real time from the video stream. A software (Video Freeze, Version 1.12.0.0, Med-Associates) performed real-time video recordings (30 frames per second) based on a threshold level (20 arbitrary units of movement), previously set and calibrated with the experimenters’ freezing scores (r^2^ = .997).

The training context (context A) was characterized by a grid floor composed of 20 stainless steel rods (diameter: 4.8mm), top and front walls made of transparent polycarbonate, a back wall made of white acrylic, stainless-steel sidewalls and drop pan. The light in the conditioning box remained on and a background noise was emitted during the training and test sessions. Context A was cleaned with alcohol 10% before and after each session.

The novel context (Context B) consisted of the conditioning chamber and stainless-steel drop pan personalized with a grid floor (20 interleaved rods of either 4.8 or 9.5mm of diameter) and white Plexiglas curved sidewalls extending across the back wall. The light in the box remained turned off and no white noise was emitted during the test. Context B was cleaned with a 5% acetic acid solution before and after each session. The dissimilarities between Contexts A and B were chosen according to previous studies that related tactile and olfactory stimuli as the most salient ones in discriminatory fear conditioning tasks (Fanselow, 1980; Huckleberry et al., 2016).

### Behavioral procedures

Figure 1 shows an overview of the experimental procedure. In all experiments, rats were randomly assigned to one of four experimental groups, balanced for mean body weight. Prior to the CFC training, rats were handled for 3 days, for 3 min each, in a room adjacent to the experimentation room. Rats were habituated to the adjacent room for at least 90 minutes before both training and testing.

**Figure 1:**
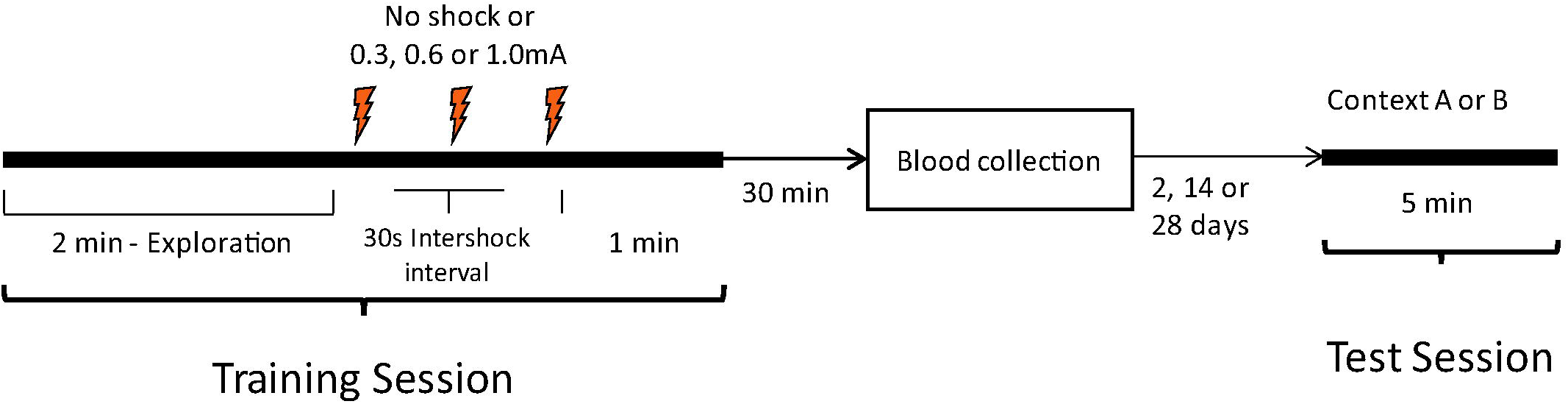
Schematic representation of the experimental design.

Two groups of animals were used as controls for this experimental design: the home-cage and the 0.0mA. Rats in the home-cage group were subjected to handling, stayed in the adjacent room with the other animals, had their blood sample collected but did not undergo training. Animals in the 0.0mA group did not receive any footshocks during the training session. On Day 1 (training session), rats (except the ones in the home-cage group) were individually transported from the adjacent room to the conditioning room and placed in context A. After 2 minutes of free exploration, three footshocks (0.3, 0.6 or 1.0mA, according to their assigned group, 1s each) were delivered with intervals of 30s between them. One minute after the third shock, the animal was removed from the apparatus and returned to its home-cage and transported back to the adjoining room. Recent or remote memory tests were performed 2, 14 or 28 days after training. For each shock intensity group, half of the animals was exposed to context A and the other half to context B. Each test lasted 5 minutes and no shock was delivered.

### Blood collection and plasma corticosterone quantification

Thirty minutes after the training, 500μL of blood was collected from each rat through the tail-clip method (Kim et al., 2018). We first cleaned the rat’s tail with lidocaine (anesthetic) to clip its most distal portion (roughly 0.5mm). The tail was carefully manipulated by the experimenter so that droplets of blood accumulated at its tip and were collected using EDTA-covered tubes. The tail tip was then treated with a balm of ketoprofen (anti-inflammatory and analgesic) and rifampicin (antibiotic).

The blood samples were centrifuged at 2300rpm for 20 min at 4°C. The extracted plasma was kept at −20°C until determination of hormone concentrations. Plasma CORT was measured using ELISA (enzyme-linked immunosorbent assay), which allows detection of specific antibodies in blood plasma using the Corticosterone Enzyme Immunoassay Kit (Arbor Assays LLC, MI, USA). This kit is supplied with clear plastic microtiter plates coated with donkey anti-sheep IgG, sheep polyclonal antibody specific for CORT, a vial of CORT at 100,000 pg/mL to be used as a standard, and the CORT-peroxidase conjugate. The ELISA procedure was conducted according to the manufacturer’s instructions. All samples were analyzed in duplicates. Later optical densities from the samples were analyzed using the Epoch Spectrophotometer system (BioTek Instruments, Inc.). CORT sample concentrations were calculated using the Arbor Assays DetectX^®^ Corticosterone (OD) software. According to the protocol in the ELISA kit, the sensitivity of the essay was determined as 18.6 pg/mL, the intra-assay coefficient of variation was in average less than 10% and the inter-assay coefficient of variation was less than 15%. ELISA standard curves were also adequate (r^2^ = 0.955 - 0.995).

### Statistical analysis

The conditioned fear response to context was quantified as the percent time the animal spent freezing in context A, whereas generalization was considered as the percent time of freezing in context B during the recall test. Behavioral results are expressed as the group mean percent freezing time ± standard error of the mean (S.E.M). Post-training plasma CORT levels are expressed as the group mean ± S.E.M.

To satisfy the requirements for the use of the ANOVA, the total freezing times and plasma CORT levels were transformed using the square root transform function (Lix et al., 1996; McDonald, 2009; Tukey, 1957) to improve data homogeneity and normality, as ascertained by Kolmogorov-Smirnov and Levene tests. The Grubbs test for outliers was used to determine extreme outliers. The transformed data was then analyzed with either a one or two-ways ANOVAs. The Student-Newmann-Keuls (SNK) *post hoc* test was further used to identify significant differences when applicable. Significance was set at *p* < 0.05. For the replication experiment, the total freezing times and plasma CORT levels were analyzed using a Student t-test for 2 independent samples, comparing means between no-shock and 0.6mA groups (CORT) or between 0.6mA groups exposed to A or B (Behavior). The effect sizes (Cohen’s d or ω2) were reported when the parametric test was found significant (“d” or “ω^2^” values above 0.8 and 0.14, respectively, are considered large effects; values between 0.5 and 0.8 (“d”) or 0.06 and 0.14 (“ω^2^) are considered moderate; and below 0.5 or 0.06, small). Significance was set at *p* < 0.05.

Associations between post-training plasma CORT and total freezing time were analyzed using Spearman product-moment correlational coefficients. Significance was set at *p* < 0.05. Linear and quadratic regression algorithms were also tested. We chose *a priori* to only test the hierarchical regression for the groups that showed significant Spearman correlations between the variables. The quadratic function (Y’ = a + b_1_X_1_ + b_2_X_1_^2^) is a second order polynomial regression representing the inverted U-shape model, which describes a parabola where Y’ is the expected CORT level and X_1_ is the observed freezing time. Higher order regressions have not been tested. Regressions coefficients of the quadratic models were considered only if the regression analysis of variance (ANOVA) was significant at p < 0.05. The model which explained most of the variance (change in r^2^ > 0.07) is given in the figures and was performed according to previous studies in the literature that investigate the relationship between memory performance and endogenous CORT levels (Lubec and Korz, 2016; McCullough et al., 2015).

## RESULTS

### CFC training and post-training plasma corticosterone levels

During training, animals that were exposed to more intense footshocks displayed more freezing on the last minute of the session (after the footshocks). This result was confirmed by a 1-way ANOVA that indicated a significant group effect [*F*_(3, 183)_ = 151.18, *p* < 0.01, ω^2^ = 0.71, Fig. 2(A)]. The SNK *post-hoc* tests indicate that all trained groups showed greater freezing levels when compared to the no-shock group (*p* < 0.01). In addition, the 0.3mA group expressed less freezing compared to the animals in groups 0.6 (*p* < 0.05) and 1.0mA (*p* < 0.01, Fig. 2(A)).

**Figure 2:**
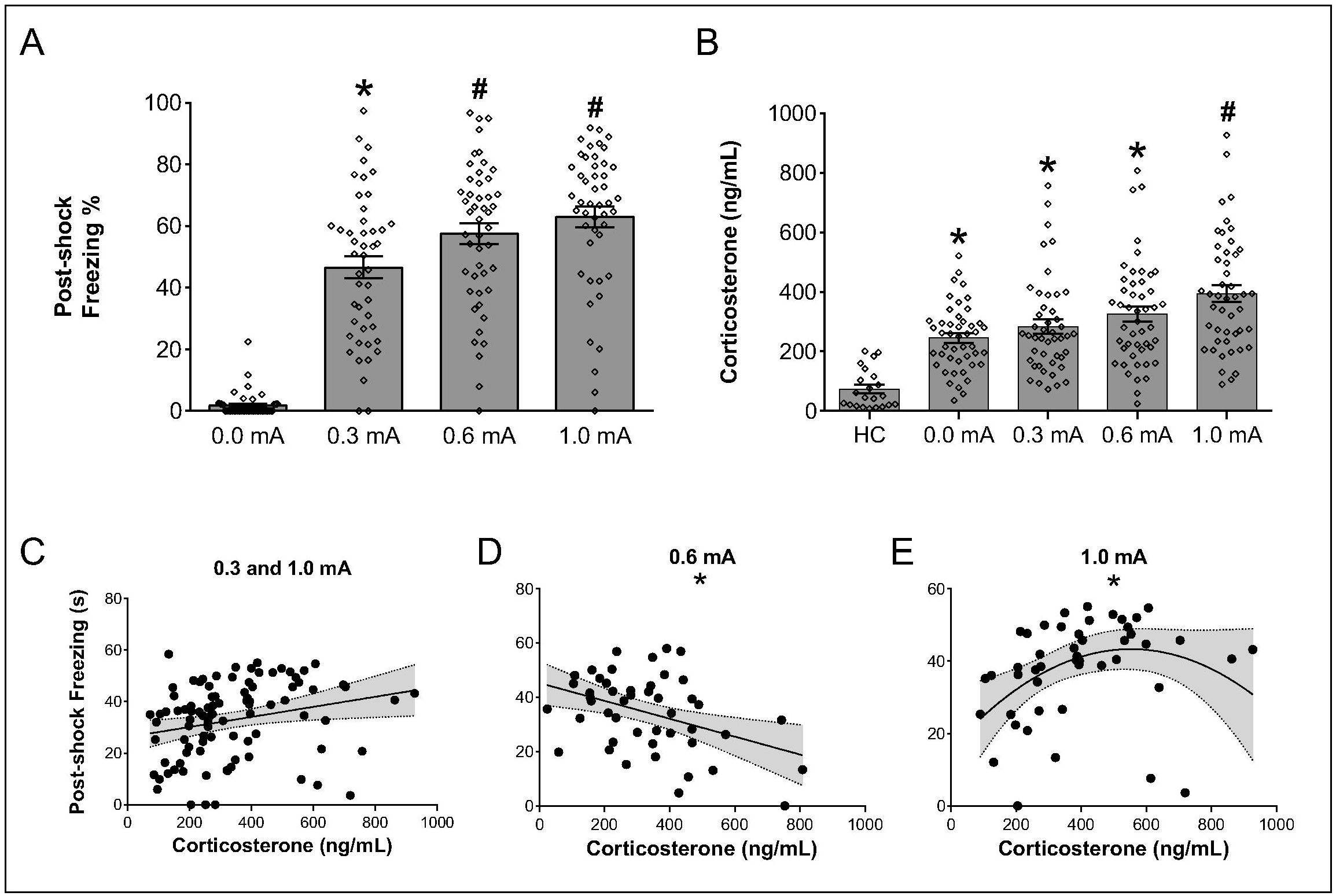
Footshock intensity during CFC training is associated with post-footshock freezing levels and post-training plasma CORT. (A) Freezing time percentage (mean ± S.E.M) during the last minute of the CFC training session. N = 46-48 by group, symbols show data from each rat. (*) p < 0.05 when compared to the 0.0mA group. (#) p < 0.01 when compared to the 0.3 and 0.0mA groups. (B) Post-training Corticosterone levels (mean ± S.E.M). (*) p < 0.0001 when compared to the Home-cage group. (#) p < 0.01 when compared to the 0.0 and 0.3mA groups. (C, D, E) Correlation between post-footshock freezing time (s) and post-training plasma CORT (ng/mL) of trained rats (0.3, 0.6 and 1.0mA groups). Animals trained with 0.6mA footshocks show a significant negative correlation (r=-.35, p = 0.02) whereas animals from group 1.0mA show a significant inverted-U curve relationship (F_(1,44)_ = 5.00, p= 0.03, r^2^ = 0.15).

Figure 2 (B) shows the post-training CORT levels. One-way ANOVA showed a significant effect [*F*_(4, 204)_ = 26.56, *p* < 0.01, ω^2^ = 0.33]. The *post-hoc* tests indicated that all groups exposed to the conditioning chamber showed higher plasma CORT levels when compared to the homecage group (*p* < 0.01). Moreover, the 1.0mA group had higher CORT levels compared to the 0.3 (*p* < 0.01) and no-shock (*p* < 0.01) groups, and the 0.6mA group was not significantly different from the no-shock (*p* = 0.09), 0.3mA (*p* = 0.26) or 1.0mA groups (*p* = 0.07).

The post-footshocks freezing time and post-training plasma CORT levels from each animal were analyzed for bivariate correlations using the Spearman test. The analysis showed a negative correlation when using the data from the 0.6mA group [*r*(46) = −.35, *p* =0.02, r^2^ = 0.12], whereas data from the 1.0mA group showed a positive correlation [*r*(45) = .41, *p* =0.00, r^2^ = 0.17]. When tested together, data from the 0.3mA and 1.0mA groups showed a positive correlation [*r*(91) = .34, p=0.01, r^2^ = 0.11]. Table 1 shows the correlation results. In each set, the Grubbs test revealed no outlier data for either freezing time or post-training plasma CORT. The analysis revealed a significant linear relationship between post-shocks freezing and plasma CORT in the 0.6mA group [*F* _(1, 45)_ = 2.23, *p* = 0.14, *r*^2^=0.17] that fits the data better than the quadratic model [*r*^2^ = 0.21, change in *r*^2^ = 0.04]. For the 1.0mA group, on the other hand, the analysis revealed a significant quadratic relationship between the post-training plasma CORT levels and post-shocks freezing time [*F*_(1, 44)_ = 5.00, *p* = 0.03, *r*^2^ = 0.15] that fits the data significantly better than the linear model [*r*^2^ = 0.06, change in *r*^2^ = 0.09]. Finally, for the combination of the 0.3 and 1.0mA groups, the analysis revealed a weak linear relationship [*F*_(1,90)_ = 3.40, *p* = 0.07, *r*^2^ = 0.06] that fits the data better than the quadratic model [*r*^2^ = 0.09, change in *r*^2^ = 0.03]. Fig. 2(C, D, E) shows the linear and quadratic fits for these groups.

**Table 1.** Spearman correlations according to footshock intensity

| Spearman correlations according to footshock intensity |  |  |  |
| --- | --- | --- | --- |
| Group | Correlation | p-value | N |
| Trained rats (0.3, 0.6, 1.0) | 0.12 | 0.16 | 141 |
| **0.3 and 1.0** | **0.34** | **0** | **93** |
| 0.3 and 0.6 | -0.1 | 0.32 | 94 |
| 0.6 and 1.0 | -0.08 | 0.47 | 95 |
| 0.3 | 0.09 | 0.57 | 46 |
| **0.6** | **-0.35** | **0.02** | **48** |
| **1.0** | **0.41** | **0** | **47** |

Note. Data in bold are significant Spearman correlations.

### CFC specificity tests

Two days after training, each group was either re-exposed to the training context (A) or exposed to a novel context (B). The Grubbs test did not find any outliers across groups. The 2-way ANOVA (Footshock × Context) showed a significant effect for Footshock [F (3, 55) = 38.35, *p* < 0.01, ω^2^= 0.56], Context [F (1, 55) = 16,04, *p* < 0.01, ω^2^= 0.08] and the interaction between Footshock and Context [F (3, 55) = 3.99, *p* < 0.01, ω^2^= 0.05]. The *post-hoc* test revealed that all groups of trained animals displayed higher freezing times when compared to the no-shock control group (0.0mA), in both contexts (p < 0.05). For animals tested in context A, the 0.3mA group presented lower freezing times than the 0.6mA and 1.0mA groups (p < 0.01), whereas there was no significant difference between these last two groups (p = 0.50). For animals tested in context B, there was no significant difference in freezing times between the different trained groups (p > 0.1). The SNK test also showed that non-trained animals and the 0.3mA groups tested in context A had freezing times similar to their counterparts tested in context B (p = 0.82 and p = 0.37, respectively), whereas animals from the 0.6 and 1.0mA groups exposed to context A showed higher freezing times compared to their counterparts exposed to context B (p < 0.01, Fig. 3(A)).

**Figure 3:**
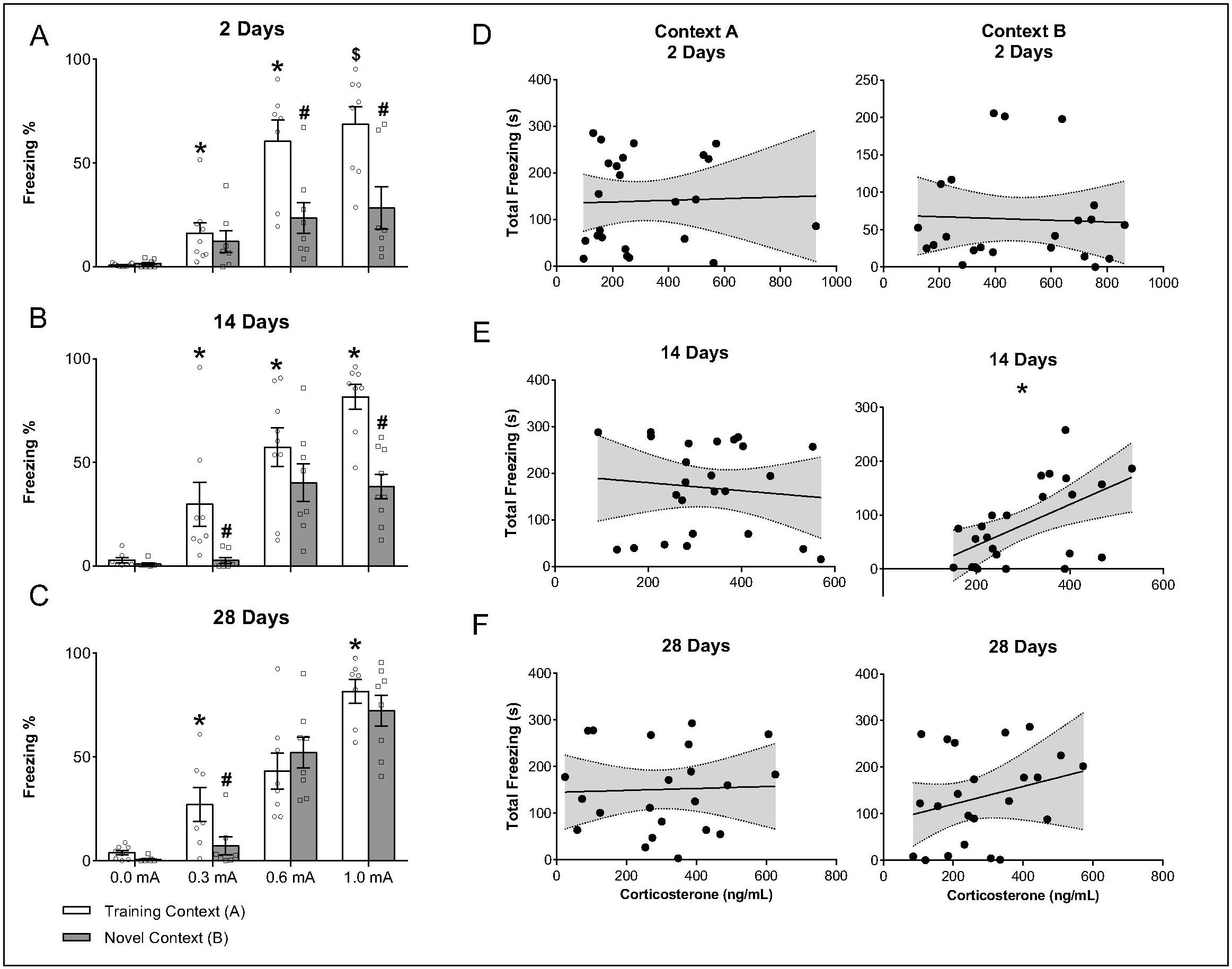
Memory retrieval tests at different timepoints. (A, B, C) Freezing time percentage (mean ± S.E.M) in context A or B during retrieval tests performed at 2, 14 or 28 days after training, respectively. N = 7-9 by group, symbols show data from each rat. (*) p < 0.05 when compared to rats trained in all lower footshock intensities and no shock. (#) p < 0.05 when compared to the group trained with the same footshock intensity but tested in context A. ($) p < 0.05 when compared to the no shock and 0.3mA groups exposed to context A. (D, E, F) Correlations between post-training plasma CORT (ng/mL) and total freezing time (s) of all trained rats (0.3, 0.6 and 1.0mA groups) tested in context A or B at 2, 14 or 28 days after training, respectively. A significant positive correlation was found only between animals that were exposed to the Context B14 days after training (r=.5, p = 0.01).

Another set of animals was tested 14 days after training. At this time-point, the two-way ANOVA showed a significant effect of Footshock [*F*_(3, 56)_ = 51.42, *p* < 0.01, ω^2^ = 0.61], Context [*F*_(1, 56)_ = 24.13, *p* < 0.01, ω^2^ = 0.11], and interaction between Footshock and Context [*F*_(3, 56)_ = 2.92, *p* = 0.04, ω^2^ = 0.02]. For animals tested in context A, the post-hoc test revealed that all trained animals had higher freezing times when compared to the no-shock group (*p* < 0.01). Moreover, the trained groups showed a training intensity-response curve, where animals trained with 1.0 mA had higher freezing than those trained with 0.6 mA (*p* < 0.05) and 0.3 mA (*p* < 0.01) and rats trained with 0.6 mA had higher freezing compared to the 0.3mA group (*p* < 0.05). Among animals exposed to context B, only those trained with 0.6 or 1.0 mA had higher freezing than the non-trained animals (*p* < 0.01, Fig. 4). The SNK test also showed that animals in groups 0.0 and 0.6mA presented similar freezing times in either contexts (A or B, *p* = 0.64 and *p* = 0.14, respectively), whereas animals from the 0.3 and 1.0mA groups, exposed to context A had higher freezing time compared to their counterparts exposed to context B (*p* < 0.01, Fig. 3(B)).

**Figure 4:**
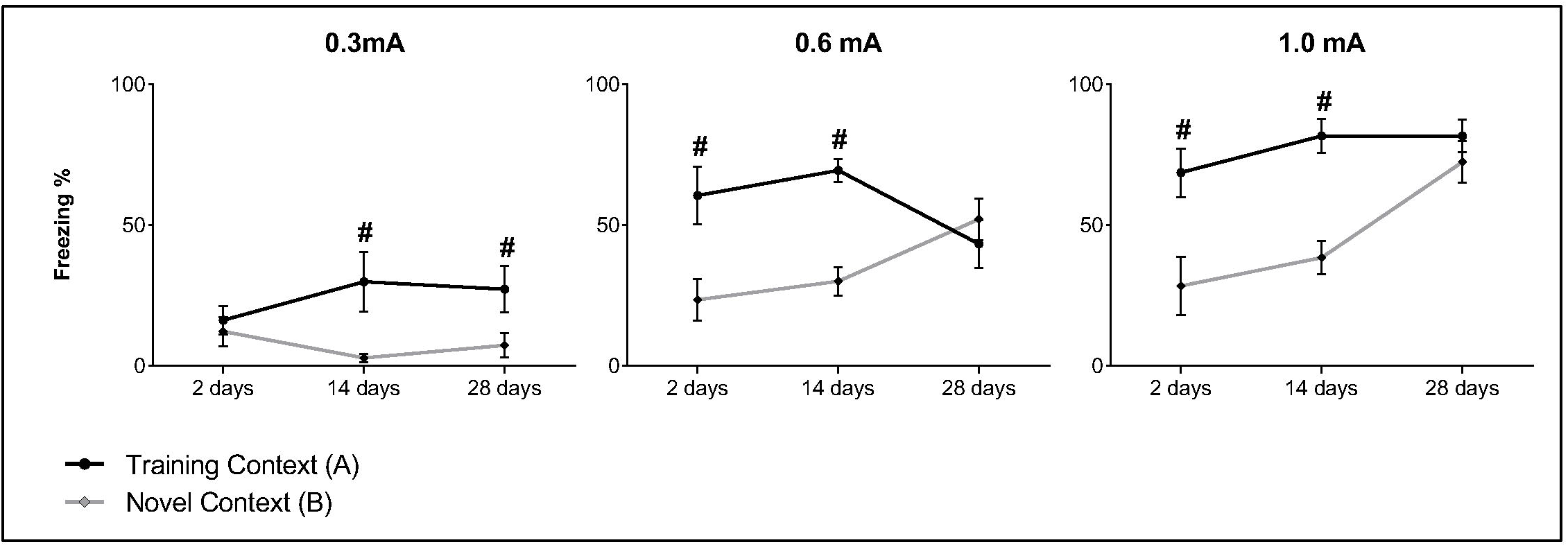
Temporal dynamics of time-dependent memory generalization. Each graph shows different groups of animals tested 2, 14 or 28 days after CFC training Rats trained with 0.3mA footshocks present preserved memory specificity at remote timepoints, whereas those trained with 0.6 or 1.0mA show generalized fear responses across contexts at 28 days. Data used for the 0.6mA, 14 days groups are from the replication experiment. (#) p < 0.05 when comparing mean freezing times of groups exposed to A or B.

The last set of animals was tested 28 days after training. The two-way ANOVA showed a significant effect of Footshock [*F*_(3, 52)_ = 76.62, *p* < 0.01, ω^2^ = 0.76], Context [*F*_(1, 52)_ = 6.21, *p* = 0.02, ω^2^ = 0.02], and interaction between Footshock and Context [*F*_(3, 52)_ = 3.36, *p* = 0.03, ω^2^ = 0.02]. The post-hoc test showed that, for animals tested in context A, all trained animals had higher freezing when compared to the no-shock group (*p* < 0.01) and the trained groups also displayed an intensity-response curve, where the 1.0mA group showed higher freezing time than the other two groups (p < 0.05), although the 0.6mA group did not differ from group 0.3mA (*p* = 0.42). For animals exposed to context B, only those trained with 0.6 or 1.0 mA showed higher freezing times than the no-shock group (*p* < 0.01, Fig. 4). The SNK test also showed that animals trained with 0.6 or 1.0 mA presented similar freezing times in both contexts (*p* = 0.36 and *p* = 0.46, respectively), whereas the 0.3mA group exposed to context A showed higher freezing time compared to its counterpart exposed to context B (*p* < 0.01, Fig. 3-C).

The post-training plasma CORT levels and freezing times during CFC test from each animal were analyzed for bivariate correlations using the Spearman test. All groups were tested together or separately, in each time point and context to which animals were exposed to (Table 2). When analyzing data from animals tested in context B, 14 days after training, there is a positive correlation between CORT and freezing times [*r*(23) = .50, *p* =0.01, r^2^ = 0.25]. As performed earlier with the training data, we analyzed non-linear interactions for the test data using the same hierarchical regression to evaluate whether data followed a linear or quadratic fit. In this set of animals, the Grubbs test for outliers revealed one outlier for CORT [*G*= 2.88, *p*<0.05], which was removed from the hierarchical analysis. This analysis revealed a linear relationship between total freezing time and post-training plasma CORT for the 14 days, Context B group [*F*_(1,21)_ = 0.56, *p* = 0.46, *r*^2^ = 0.30] that fit the data significantly better than the quadratic model [*r*^2^ = 0.32, change in *r*^2^ = 0.02]. Fig. 3 (D, E, F) shows the correlation and linear fits for all trained groups exposed to context A or B.

**Table 2.**
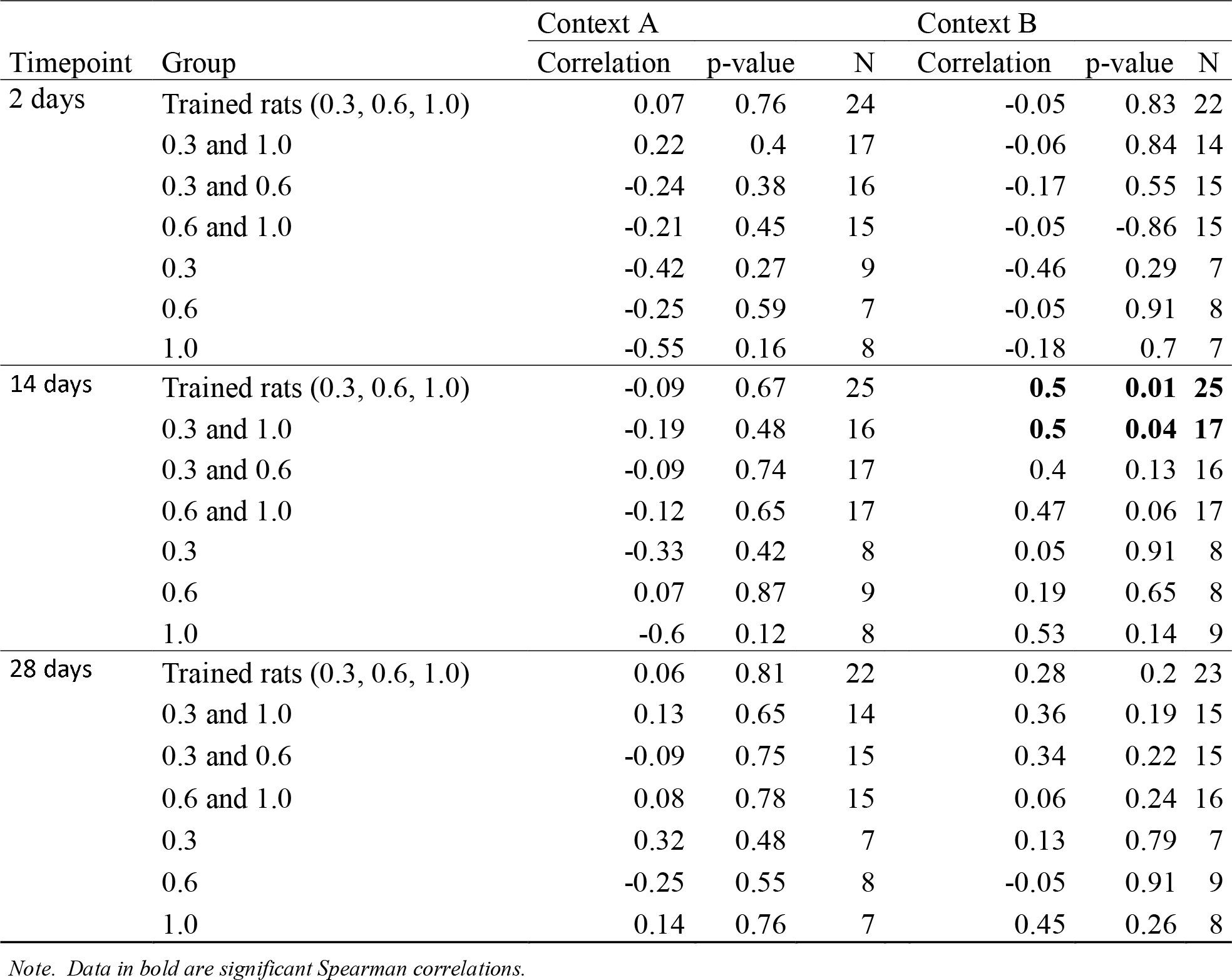
Spearman correlations according to shock intensity, timepoint and test context

| Timepoint | Group | Context A |  |  | Context B |  |  |
| --- | --- | --- | --- | --- | --- | --- | --- |
|  |  | Correlation | p-value | N | Correlation | p-value | N |
| 2 days | Trained rats (0.3, 0.6, 1.0) | 0.07 | 0.76 | 24 | -0.05 | 0.83 | 22 |
|  | 0.3 and 1.0 | 0.22 | 0.4 | 17 | -0.06 | 0.84 | 14 |
|  | 0.3 and 0.6 | -0.24 | 0.38 | 16 | -0.17 | 0.55 | 15 |
|  | 0.6 and 1.0 | -0.21 | 0.45 | 15 | -0.05 | -0.86 | 15 |
|  | 0.3 | -0.42 | 0.27 | 9 | -0.46 | 0.29 | 7 |
|  | 0.6 | -0.25 | 0.59 | 7 | -0.05 | 0.91 | 8 |
|  | 1.0 | -0.55 | 0.16 | 8 | -0.18 | 0.7 | 7 |
| 14 days | Trained rats (0.3, 0.6, 1.0) | -0.09 | 0.67 | 25 | **0.5** | **0.01** | **25** |
|  | 0.3 and 1.0 | -0.19 | 0.48 | 16 | **0.5** | **0.04** | **17** |
|  | 0.3 and 0.6 | -0.09 | 0.74 | 17 | 0.4 | 0.13 | 16 |
|  | 0.6 and 1.0 | -0.12 | 0.65 | 17 | 0.47 | 0.06 | 17 |
|  | 0.3 | -0.33 | 0.42 | 8 | 0.05 | 0.91 | 8 |
|  | 0.6 | 0.07 | 0.87 | 9 | 0.19 | 0.65 | 8 |
|  | 1.0 | -0.6 | 0.12 | 8 | 0.53 | 0.14 | 9 |
| 28 days | Trained rats (0.3, 0.6, 1.0) | 0.06 | 0.81 | 22 | 0.28 | 0.2 | 23 |
|  | 0.3 and 1.0 | 0.13 | 0.65 | 14 | 0.36 | 0.19 | 15 |
|  | 0.3 and 0.6 | -0.09 | 0.75 | 15 | 0.34 | 0.22 | 15 |
|  | 0.6 and 1.0 | 0.08 | 0.78 | 15 | 0.06 | 0.24 | 16 |
|  | 0.3 | 0.32 | 0.48 | 7 | 0.13 | 0.79 | 7 |
|  | 0.6 | -0.25 | 0.55 | 8 | -0.05 | 0.91 | 9 |
|  | 1.0 | 0.14 | 0.76 | 7 | 0.45 | 0.26 | 8 |

*Note. Data in bold are significant Spearman correlations.***Replication of the 0.6mA group (moderate CFC)**

### Replication of the 0.6mA group (moderate CFC)

Due to the large variability of the 0.6mA groups exposed to context A or B at 14 days post-training, this experiment was replicated with a new set of animals (N= 23), tested 14 days later, when half was exposed to context A, and the other half to context B. A no-shock group of 9 animals was used as a control for the post-training plasma CORT levels. The Student t-test showed a significant increase in plasma CORT levels for the 0.6mA group [*t*(30) = −2.39, *p* < 0.05, *d* = −0.94]. Moreover, animals trained with 0.6mA and tested in context A presented higher freezing times than their counterparts tested in context B [*t*(21) = 6.04, *p* < 0.01, *d* = 2.47), Table 3].

**Table 3.** Post-training plasma CORT levels of the replication experiment

| Group | Corticosterone ± S.E.M<br>(ng/mL) | N |
| --- | --- | --- |
| 0.0mA | 128.5 ± 21.24 | 9 |
| 0.6mA | 232.2 ± 29.64* | 24 |

| Freezing times during the retrieval tests at 14 days after the training |  |  |
| --- | --- | --- |
| Group | Freezing time ± S.E.M (%) | N |
| 0.6mA Context A | 69.41 ± 4.08 | 12 |
| 0.6mA Context B | 29.97 ± 5.10* | 12 |

Note. (\*) $p < 0.05$ when compared to the control groups

### Memory generalization rate

Figure 4 shows mean freezing times from all groups tested 2, 14 or 28 days after training (the results from the replication of the 14 days’ 0.6mA group was used instead of the original 0.6mA group). Animals exposed to context A show a trend of high freezing times at all timepoints. Animals exposed to context B, however, show a trend of increased freezing times as memory ages – except for animals trained with 0.3mA footshock. Even though inferential statistical tests between time points were not performed due to the fact that the sets of animals tested in different time intervals were trained separately, it is possible to infer memory transformation when analyzing *post-hoc* results. For the 0.3mA group, exposure to contexts A and B induced similar freezing times on day 2 (*p* = 0.37), whereas on days 14 and 28 freezing was higher in context A (*p* < 0.01). On the other hand, groups 0.6 or 1.0mA showed higher freezing times in context A, compared to context B at 2 and 14 days, but similar freezing (0.6mA: *p* = 0.13, 1.0mA: *p* = 0.46) times between contexts at 28 days after training.

## DISCUSSION

Our findings indicate that the intensity of the footshock in the CFC training is positively associated with the specificity and rate of generalization of fear memories. This effect may be due, although not entirely dependent, to post-training plasma CORT (Kaouane et al., 2012; Pedraza et al., 2016). Our results partially replicate those from Cordero and colleagues (1998), which suggested a linear relationship between post-training plasma CORT levels and freezing times during CFC training (Cordero et al., 1998). Our data, however, only points to a linear relationship between CORT and freezing if we ignore the animals that underwent a moderate intensity training (0.6mA group). Indeed, when we restrict our analysis to individual subsets (0.6 or 1.0mA), the relationship found between freezing times and plasma CORT follows different trendlines (i.e. negative linear and inverted u-curve, respectively). Our evidence of a non-linear interaction between these variables is strengthened by the study of Marchand and colleagues (2007), whose results did not corroborate a positive linear correlation between freezing levels during the training session and the post-training plasma CORT levels (Marchand et al., 2007). We should note, however, that their study did not contemplate the analysis of quadratic interactions between the variables. Furthermore, our results show that, on average, rats that released moderate amounts of CORT displayed greater freezing times during the last minute of the CFC training, whereas lower or higher CORT levels were associated to lower post-shock freezing times. Hence, it is possible to speculate that the relationship between CORT and training intensity follows a more complex trend than a simple linear relationship.

We also found that increasing training intensity elicits higher freezing times during retrieval tests when animals are re-exposed to the training context even at remote timepoints, which corroborate the findings from previous studies (Cordero et al., 1998; Kaouane et al., 2012; Pedraza et al., 2016; Poulos et al., 2016). Norepinephrine (Atucha et al., 2017; Gazarini et al., 2015, 2013; Gold and Van Buskirk, 1976, 1975) and CORT (Abrari et al., 2009; Kaouane et al., 2012) have been assumed to enhance memory consolidation, following an inverted U-curve dose-effect relation. It is possible, therefore, that stronger CFC protocols induce stronger activation from the hypothalamus-pituitary-adrenal (HPA) axis and noradrenergic system. Moreover, our data show that the temporal rate of generalization can be modulated just by increasing footshock intensity during training, without changing the number of footshocks. This result is in line with the study of Pedraza and colleagues (2016), who reported that training protocols with more and stronger footshocks lead to early memory generalization. Curiously, however, rats in our study that underwent the lowest intensity training protocol (0.3mA) presented similar freezing times (albeit significantly different from the no-shock groups) in contexts A and B, when tested 2 days after training. This could reflect a sensitization to all novel situations due to the recentness of the training session, instead of an error of retrieval (Richardson, 2000). This alternative explanation to recent memory generalization is supported by the fear responses of rats trained with 0.3mA footshocks and tested after 14 or 28 days, that shows discrimination between contexts A and B. Moreover, rats exposed to context B at the remote timepoints presented similar freezing times to the no-shock groups tested in the same context.

It is worth noting that the only significant linear relationship between freezing times and post-training plasma CORT occurred at 14 days (post training) in animals exposed to context B. According to the literature, this is the earliest time point which animals tend to generalize contextual fear responses (Kim and Fanselow, 1992; Pedraza et al., 2016; Wiltgen and Silva, 2007). Thus, our results suggest that the process of time-dependent generalization of fear in novel contexts begins or is underway about this time point, and it is characterized by a linear relationship to the post-training plasma CORT levels (i.e. lower CORT levels elicit low freezing times in context B, whereas higher CORT levels elicit higher freezing times).

According to the systems consolidation theory, memories change dependence from hippocampus to neocortical areas overtime, which is usually associated with time-dependent generalization (Biedenkapp and Rudy, 2007; Frankland et al., 2004; Wang et al., 2009; Wiltgen et al., 2010; Wiltgen and Silva, 2007). The rate by which this transfer occurs is thought to be modulated by the intensity of the training protocol and possibly, according to our results, by the levels of plasma CORT released afterwards. This modulatory role of CORT in memory generalization is probably due to simultaneous but different processes happening in the prefrontal cortex and hippocampus (i.e. complementary learning systems – McClelland et al., 1995; Norman, 2010; O’Reilly et al., 2014; O’Reilly and Norman, 2002).

Previous studies have shown that the difference in the ratio of activation of MR and GRs is fundamental for memory consolidation (Cordero and Sandi, 1998; Pugh et al., 1997a, 1997b; Sandi, 1998). It has long been established that the prefrontal cortex shows 25% less GRs when compared to the hippocampus (Meaney and Aitken, 1985), and a high density of MRs (Reul and Kloet, 1985), whereas the hippocampus and amygdala show high density of GRs and few MRs (Van Eekelen et al., 1988). Consequently, the same levels of plasma CORT lead to different ratios of activation of both GR and MR in these regions (Meijer et al., 2010; for a review on the subject, see De Kloet, 2013). McCullough and colleagues (2015) have suggested that, due to the difference of GR / MR ratios in these regions, memory performance for contextual (hippocampus dependent) and other memory tasks (neocortex dependent) follows two separate U-inverted curves, associated to increasing plasma CORT levels (McCullough et al., 2015). Therefore, it is possible to speculate that, after a low intensity CFC training, low concentrations of plasma CORT were able to successfully reach the peak of hippocampal GR/MR activation and, therefore, optimum synaptic plasticity in this area but not in the neocortex. This may explain why our lower intensity CFC training retains memory specificity for longer periods of time (see Fig. 5). Moreover, higher concentrations of CORT are required to reach both optimal hippocampal and neocortical GR/MR ratio peak (for a review, see Van Ast et al., 2014). In a strong intensity CFC training, we hypothesize that the high plasma CORT levels might rise beyond optimal hippocampal activation, ascribing non-specific contextual features to the memory trace which, in turn, could lead to subsequent retrieval errors (Hunsaker and Kesner, 2013, 2008; Kesner and Hunsaker, 2010; Rudy and O’Reilly, 1999). This could explain the fact that high CORT levels during memory consolidation resulted in generalized, remote memories.

**Figure 5:**
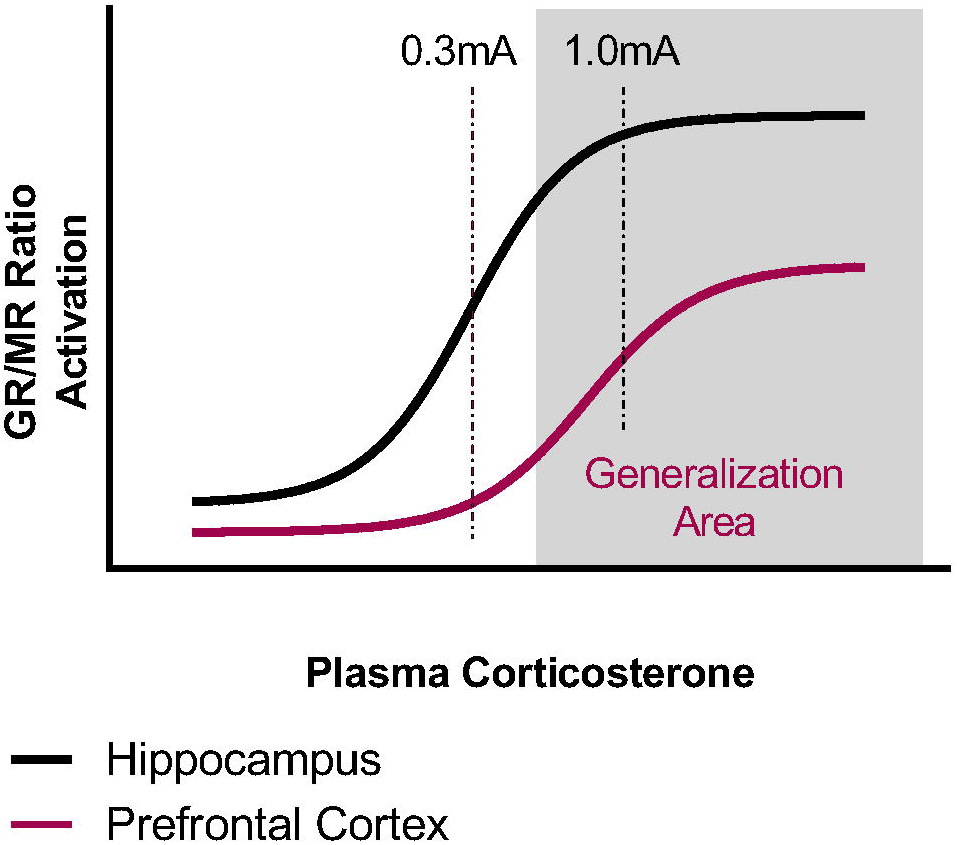
A graphical representation on the hypothesis that simultaneous corticosterone (CORT) receptors activation in hippocampus and prefrontal cortex in early consolidation elicits time-dependent generalization. Glucocorticoid Receptors (GR) / Mineralocorticoid Receptors (MR) density ratios are different in the hippocampus and prefrontal cortex, and can be represented by two different sigmoid curves that describes GR/MR activation for each region. Higher CORT levels, associated with high intensity trainings, may be required for optimal hippocampal and neocortical GR/MR activation, enabling time-dependent memory generalization. Lower CORT levels elicit optimal hippocampal but not neocortical GR/MR activation, hence inducing slower systems consolidation.

Many previous studies have successfully tested memory discrimination by presenting different groups of rats to either context A or B (Baldi et al., 2004; Biedenkapp and Rudy, 2007; Haubrich et al., 2016; Pedraza et al., 2016; Poulos et al., 2016; Wiltgen and Silva, 2007). A possible limitation for this protocol, however, is that rats were never tested in both contexts. Presenting animals to both contexts in a counterbalanced order could be a more direct way to evaluate contextual memory specificity (Bueno et al., 2017; Huckleberry et al., 2016; Poulos et al., 2016; Wang et al., 2009). Nevertheless, our protocol was chosen to avoid any possible influence of memory reactivation induced by reexposure to the training context.

In conclusion, our study sheds light on a possible role for CORT in modulating time-dependent memory generalization, probably associated with the process of systems consolidation. Plasma CORT levels and freezing time elicited by the CFC training seem to follow a more complex relationship than the previously thought linear correlation. In addition, our findings support the hypothesis that the CFC training intensity modulates the rate of generalization, with higher intensities eliciting faster generalization and lower intensities prolonging memory specificity. Finally, approximately 14 days after training may be a tipping time point for memory transformation modulated by post-training plasma CORT levels.

## ACKNOWLEDGEMENTS

MdSC elaborated and executed experiments, analyzed data and wrote this article; BdSV and GDVG executed experiments; JPQP analyzed data and wrote this article; PAT and RVF supervised experiments and wrote this article.

